# Nicotinamide Deficiency in Primary Open-Angle Glaucoma

**DOI:** 10.1101/571638

**Authors:** Judith Kouassi Nzoughet, Juan Manuel Chao de la Barca, Khadidja Guehlouz, Stéphanie Leruez, Laurent Coulbault, Stéphane Allouche, Cinzia Bocca, Jeanne Muller, Patrizia Amati-Bonneau, Philippe Gohier, Dominique Bonneau, Gilles Simard, Dan Milea, Guy Lenaers, Vincent Procaccio, Pascal Reynier

**Affiliations:** Equipe Mitolab, Unité Mixte de Recherche MITOVASC, CNRS 6015, INSERM U1083, Université d’Angers, Angers, France; Département de Biochimie et Génétique, Centre Hospitalier Universitaire, Angers, France; Département d’Ophtalmologie, Centre Hospitalier Universitaire, Angers, France; Service de Biochimie, EA4650, Centre Hospitalier Universitaire, Caen, France; Singapore Eye Research Institute, Singapore National Eye Centre, Duke-NUS, Singapore

**Keywords:** Primary open-angle glaucoma, Metabolomics, Mitochondrial dysfunction, Nicotinamide, Vitamin B3, Nicotinamide adenine dinucleotide, NAD, Optic nerve, Optic neuropathy

## Abstract

**Purpose:** To investigate the plasma concentration of nicotinamide in primary open-angle glaucoma (POAG).

**Methods:** Plasma of 34 POAG individuals were compared to that of 30 age-and sex-matched controls using a semi-quantitative method based on liquid chromatography coupled to high-resolution mass spectrometry. Subsequently, an independent quantitative method, based on liquid chromatography coupled to mass spectrometry, was used to assess nicotinamide concentration in the plasma from the same initial cohort and from a replicative cohort of 20 POAG individuals and 15 controls.

**Results:** Using the semi-quantitative method, the plasma nicotinamide concentration was significantly lower in the initial cohort of POAG individuals compared to and further confirmed in the same cohort, using the targeted quantitative method, with mean concentrations of 0.14 µM (median: 0.12 µM; range: 0.06-0.28 µM) in the POAG group (−30 %; *p* = 0.022), and 0.19 µM (median: 0.18 µM; range: 0.08-0.47 µM) in the control group. The quantitative dosage also disclosed a significantly lower plasma nicotinamide concentration (−33 %; *p* = 0.011) in the replicative cohort with mean concentrations of 0.14 µM (median: 0.14 µM; range: 0.09-0.25 µM) in the POAG group, and 0.19 µM (median: 0.21 µM; range: 0.09-0.26 µM) in the control group.

**Conclusions:** Glaucoma is associated with lower plasmatic nicotinamide levels, compared to controls, suggesting that nicotinamide supplementation might become a future therapeutic strategy. Further studies are needed, in larger cohorts, to confirm these preliminary findings.

## INTRODUCTION

Glaucoma, the leading cause of irreversible blindness worldwide, is due to a progressive optic neuropathy involving the loss of retinal ganglion cells (RGCs) ^1^. Although age and increased intraocular pressure (IOP) are the main risk factors of the disease, other factors may contribute to the occurrence and progression of glaucoma, such as genetic variants, which account for approximately 5 % of the cases, together with vascular impairment, and metabolic disturbances^2^.

Since the local absence of myelinated axons in the intraocular portion of the optic nerve leads to high energy requirements, the question of mitochondrial dysfunction has been raised in glaucoma similarly to what is observed in hereditary optic neuropathies ^3^. Indeed, several studies have revealed a true respiratory chain deficiency in glaucoma ^4,5^. The central role of mitochondrial dysfunction was recently demonstrated in a DBA/2J mouse model of glaucoma with high IOP ^6,7^. These authors highlighted decreased retinal levels of nicotinamide adenine dinucleotide (NAD), an essential oxidation-reduction cofactor, and showed that the oral administration of high doses of nicotinamide, a precursor of NAD, structurally and functionally prevented the loss of RGCs, posing the rationale for a translational application in humans ^8^.

Nicotinamide, also known as vitamin B3 or PP (pellagra-preventive) vitamin, is a water-soluble vitamin, the deficiency of which causes pellagra, a systemic condition associating diarrhoea, dermatitis and dementia, and ultimately leading to death. Despite its potential role in the pathogenesis of glaucoma, no study to our knowledge has yet established the involvement of nicotinamide in individuals with primary open-angle glaucoma (POAG) ^9^.

To gain insight into the pathophysiology of POAG, we applied a non-targeted metabolomics approach, based on liquid chromatography coupled to high resolution mass spectrometry (LC-HRMS) ^10^, to compare the plasma of individuals with POAG and controls. This study, showing that nicotinamide was the most discriminating metabolite of the signature, led us to investigate the plasma concentration of nicotinamide in individuals with POAG, as reported here.

## METHODS

### Ethics Statement

Participants were included in the study after having given their informed written consent for the research. The study was conducted according to the ethical standards of the Helsinki Declaration and its later amendments, and with the approval of the University of Angers ethical committee (Comité de Protection des Personnes (CPP) OUEST 2), agreement number: CB 2013-04.

### Study participants

Individuals were recruited from the Department of Ophthalmology of Angers University Hospital, France. The initial diagnosis of POAG was based on consensual criteria, i.e. glaucomatous optic nerve damage with progressive optic disc cupping, associated with an IOP >21 mmHg ^11^. All the patients with POAG had an elevated IOP at the time of initial diagnosis, as well as open irido-corneal angles, as determined by gonioscopic examination. Individuals with isolated ocular hypertension, normal tension glaucoma, or any secondary form of glaucoma, were excluded from the study. Standard automated perimetry (Humphrey field analyser, Carl Zeiss, Dublin, CA, USA) with the 24-2 SITA-Fast algorithm was performed on all individuals with POAG, and values of the visual field mean defect (VF-MD) were used to grade the severity of POAG as “mild” with values lower than −6 dB, “moderate” with values between −6 dB and −12 dB, and “severe” with values higher than −12 dB (perimetric Hoddap-Parrish-Anderson criteria). The reliability indices retained were false positive or false negative rates under 15 %, and fixation losses under 20 %. The other tests performed on patients with POAG included evaluation of the thickness of the retinal nerve fibre layer (RNFL), using spectral domain optical coherence tomography (OCT), and measurement of the central corneal thickness (CCT) (Cirrus OCT, Carl Zeiss Meditec, Dublin, CA, USA). The best-corrected visual acuity was measured using the Monoyer decimal charts, with the results converted into logMAR units for statistical analysis. The IOP was measured using the Goldmann applanation tonometer. The history of glaucoma treatment was documented.

Control subjects were selected among healthy individuals undergoing cataract surgery at the same Department of Ophthalmology. Their inclusion criteria were: visual acuity ≥ 20/50 and the absence of any other associated ocular condition, excepting cataract. The exclusion criteria were: a family history of glaucoma, ocular hypertension or any other intraocular pathology, including retinal disorders.

Our study was carried out on two distinct cohorts recruited from the Department of Ophthalmology of Angers University Hospital. The first cohort, referred as the “initial cohort”, was composed of 34 individuals with POAG and 30 controls, and the second cohort, referred as the “replicative cohort”, was composed of 20 individuals with POAG and 15 controls. The initial cohort was subjected to a non-targeted metabolomics study, which led to the discovery of nicotinamide deficiency. This was followed by a quantitative analysis as developed in the Department of Biochemistry of Caen University Hospital, France. The replicative cohort was used only for the specific quantitative analysis of nicotinamide.

Blood samples from each participant were collected in heparin tubes at least three hours after the last meal. The transfer of the blood tubes was carried out according to a very strict protocol, securing the fastest possible storage at −80 degrees C. Thus, after blood sampling, the tubes were immediately transported on ice to the certified Biological Resource Center (Hospital of Angers), where they were immediately processed for centrifugation (10 minutes at 3000 g at +4 °C) to recover the supernatant (plasma), which was aliquoted in 500 microliter aliquots, and immediately stored at −80°C until further analysis. The delay between sampling and storage was less than one hour for every included subject.

### Non-targeted semi-quantitative LC-HRMS nicotinamide analysis of plasma samples from the initial cohort

The non-targeted LC-HRMS analysis was performed according to a method designed for the semi-quantitative measurement of 501 metabolites ^10^. Briefly, metabolites were extracted from plasma samples using ice-cold methanol. The extracts were analysed by reverse phase (RP) ultra-high-performance liquid chromatography (UHPLC, Dionex™ UltiMate 3000) coupled to a high-resolution mass spectrometer (HRMS, Thermo Scientific™ Q Exactive™ platform). Acquisitions were performed in heated electrospray positive ionization (HESI+) mode. The semi-quantitative measurement of nicotinamide was based on an in-house library composed of 501 endogenous metabolites, created using the Mass Spectrometry Metabolite Library of Standards (IROA Technology, Bolton, MA, USA). The method was validated over three days, and the extraction efficiency as well as the accuracy, precision, repeatability, and linearity of the method were assessed to ensure the quality of the results ^10^.

The parameters of nicotinamide in the non-targeted method were the following: ionization: positive mode; RT: 1.66 min; Formula: C_6_H_6_N_2_O; M+H: 123.0553; Fragment ions: 80.0501 and 96.0449. The repeatability (CV% performed on 6 duplicates) of the method for nicotinamide was as follow: 5.5% for peak area, 7.6% for peak intensity, 0.7% for retention time (RT) and 0% for m/z ratio. Mass spectrometry and chromatography accuracies were also satisfactory, with respectively 1 Δppm and 0.05 ΔRT; R^2^ for dilutions linearity (1, 1/2, 1/4 dilutions) was equal to 0.9.

### Quantitative LC-MS/MS nicotinamide analysis of plasma samples from the initial and replicative cohorts

A blind independent external validation of nicotinamide dosage was performed on plasma samples from both the initial and replicative cohorts using a targeted LC-MS/MS method specifically designed for the quantification of nicotinamide. Nicotinamide (NM) and its isotope-labelled analogue, nicotinamide-d_4_ (NM-d_4_), were purchased from LGC Standards GmbH (Wesel, Germany). Fifty microliters of plasma were mixed with 20 µL Internal Standard (IS) solution (NM-d_4_), and 130 µL of a cold methanol/acetonitrile solution (50/50; V/V) to precipitate proteins. Samples were incubated on ice for 5 min, and then centrifuged at 10 000 g for 5 min. Fifty µL of supernatant were mixed with 550 µL of water and filtered (0.45 μm) before injection into the chromatography and mass spectrometry system.

Liquid chromatography was conducted on a UFLC Prominence chromatographic system (Shimadzu, Kyoto, Japan) connected to a SCIEX QTRAP^®^ 5500 mass spectrometer, equipped with a turbo V ion spray source (SCIEX, Toronto, Canada). Six µL of supernatant were injected, and chromatographic separation was performed at +40 °C using a Pursuit pentafluorophenyl (PFP) column (150 x 2.1 mm, 3.5 µm; Agilent technologies, Santa Clara, CA, USA) connected to a guard column (Pursuit PFP). The flow rate was 0.4 ml.min^-1^. A gradient mobile phase was performed and started with 98 % mobile phase A (0.1% formic acid in water) and 2 % mobile phase B (methanol). After 1.5 min post-injection, the percentage of mobile phase B increased linearly from 2 % to 80 % in 1 min, and stayed at 80 % mobile phase B during 0.5 min. The return to baseline conditions (2 % B) was operated after 4 min and the system was allowed to stabilize for 2.3 min before the next injection. The total chromatographic run time was 6.3 min.

Mass spectrometry analysis was conducted using the electrospray ion (ESI) source in the positive mode. The parameters of the ion source were as follows: temperature 450 °C, ESI voltage 5500 V, Gas GS1 70 psi, Gas GS2 60 psi, CAD gas 8 psi, and Curtain gas 40 psi. For nicotinamide quantification, Multiple Reaction Monitoring (MRM) transitions were respectively m/z 123→80 and m/z 127→84 for nicotinamide and nicotinamide-d_4_ respectively. For nicotinamide transition, the instrument parameters were 91 V, 27 V, and 12 V for DP, CE, and CXP, respectively. For nicotinamide-d_4_ transition, the instrument parameters were 81 V, 27 V, and 38 V for DP, CE, and CXP, respectively.

Five standard calibration points were made in water at final concentrations of 0.082, 0.205, 0.410, 0.819, and 1.639 µM for nicotinamide. A solution of nicotinamide-d_4_ was prepared by dilution in water at a final concentration of 3.966 µM (IS solution).

Evaluation of the sensitivity and specificity of the protocol showed that the targeted LC-MS/MS method gave good results. The calibration curve was linear up to 200 µg/L (r>0.999), the limit of quantification was 5 µg/L, and the recovery rate was 101±3 % in plasma samples spiked with nicotinamide. During the reproducibility assay, the coefficients of variation (CV) were lower than 5 % at three levels of concentration (CV = 4.8%, 20.4±1.0 µg/L for the low-level control). The retention times were 1.73 min and 1.71 min for nicotinamide and nicotinamide-d_4_, respectively. Typical chromatograms for nicotinamide and nicotinamide-d_4_ in plasma samples are shown in the supplementary Figure.

### Statistical analyses

The data matrix from non-targeted metabolomics contained one hundred and sixty metabolites; univariate analysis was performed using the non-parametric Wilcoxon rank sum test with Benjamini-Hochberg correction and keeping the False Discovery Rate (FDR) below 5%. These analyses were conducted using Metaboanalyst v4.0 ^12^.

Univariate analyses of clinical data were carried out using two-tailed Student’s *t*-test, with differences being considered significant at *p* < 0.05. A median test was used to compare the median concentrations of nicotinamide found in individuals with POAG *versus* controls, in both the initial and replicative cohorts. The level of significance for the two-tailed test was set at α = 0.05. This analysis was performed using SPSS Statistics v22 (IBM, Bois-Colombes, France).

The Chi-squared test was performed to assess the independence between POAG and control, in relation to the distribution of the blood collection hour (morning *vs*. afternoon).

## RESULTS

This investigation was exclusively designed for a dedicated cohort of glaucoma patients and controls, and POAG was the only outcome under consideration.

As the literature does not report diurnal variations in vitamin B3 levels, we included patients who were selected in our ophthalmic clinics within the daily operating hours (from 8am to 4pm). In addition, subjects were included only if they had been fasting for at least 3 hours, before reaching the hospital. However, to exclude an eventual bias due to the collection time, we statistically compared the collection times of the patients and control cohorts, without finding significant heterogeneity (supplementary Table).

### Clinical characteristics of individuals with POAG and controls

Comparisons between individuals with POAG (n=34) and controls (n=30) from the initial cohort, in terms of demographic and comorbidity data, medical conditions and general ophthalmological features, are presented in Table 1. There were no significant differences between the two groups in terms of mean age, sex ratio, systemic medications, or mean IOP.

**Table 1:** Characteristics of individuals from the initial cohort. Demographic data and comorbidity status, systemic medications, ophthalmological features and glaucoma medication of individuals with POAG compared to controls. BMI: body mass index (weight/height^2^). IOP: intraocular pressure; CCT: central corneal thickness; RNFL: retinal nerve fibre layer; VF-MD: visual field mean defect.

|  | **POAG**<br>**(N=34)** | **Controls**<br>**(N=30)** | ***p*-**<br>**value** |
| --- | --- | --- | --- |
| **Demographic data and**<br>**comorbidity** |  |  |  |
| Average age (y) | 73.06 | 73.77 | 0.65 |
| Females (%) | 50 | 50 | 1 |
| Mean BMI (kg/m<sup>2</sup>) | 26.22 | 26.99 | 0.59 |
| Diabetes (%) | 17.65 | 3.33 | 0.10 |
| Hypertension (%) | 50 | 63.33 | 0.29 |
| Hyperlipidaemia (%) | 26.47 | 43.33 | 0.165 |
| Thyroid disease (%) | 11.76 | 13.33 | 0.29 |
| **Systemic medications** |  |  |  |
| Anti-hypertensives (%) | 47.06 | 63.33 | 0.19 |
| Lipid-lowering medications (%) | 23.53 | 43.33 | 0.09 |
| Antiplatelet therapy (%) | 26.47 | 36.67 | 0.39 |
| Oral diabetes medications (%) | 14.71 | 13.33 | 0.88 |
| Insulin (%) | 2.94 | 0 | 0.32 |
| Corticosteroids (%) | 2.94 | 3.33 | 0.93 |
| Thyroid hormone (%) | 17.65 | 13.33 | 0.64 |
| Oestrogen (%) | 0 | 0 | 1 |
| Vitamin D (%) | 11.76 | 20 | 0.38 |
| **Ophthalmological features and**<br>**glaucoma medication** |  |  |  |
| Mean visual acuity (LogMar) | +0.12 | +0.13 | 0.91 |
| Mean IOP (mmHg) | 13.42 | 14.10 | 0.27 |
| Mean CCT (μm) | 529.95 | - | - |
| Average RNFL thickness (μm) | 66.91 | - | - |
| Mean VF-MD (dB),<br>(eye with worse MD) | -6.83 | - | - |
| Glaucoma severity (%) |  |  |  |
| Mild | 82.35 | - | - |
| Moderate | 5.88 | - | - |
| Severe | 11.77 | - | - |
| Glaucoma medications (%) |  |  |  |
| Beta-blockers | 55.88 | - | - |
| Prostaglandin analogue | 67.65 | - | - |
| Alpha-2-agonists | 11.76 | - | - |
| Carbonic anhydrase inhibitor | 26.47 | - | - |

Comparisons between individuals with POAG (n=20) and controls (n=15) from the replicative cohort, in terms of demographic and comorbidity data, medical conditions and general ophthalmological features are presented in Table 2. There was no significant differences between the two groups in terms of mean age, sex ratio, or systemic medications, except for anti-hypertensives (*p*<0.02) and lipid-lowering medications (*p*<0.04), which were significantly lower in individuals with POAG than in controls. In contrast to the initial cohort, the replicative cohort showed a difference between the two groups regarding the IOP, which was significantly higher in POAG individuals compared to controls (p<0.001), the discrepancy with the initial cohort being related to the presence in the replicative cohort of patients with an insufficiently efficacious treatment for IOP.

**Table 2:** Characteristics of individuals from the replicative cohort. Demographic data and comorbidity status, systemic medications, ophthalmological features and glaucoma medication of individuals with POAG compared to controls. BMI: Body mass index (weight/height^2^). IOP: intraocular pressure; CCT: central corneal thickness; RNFL: retinal nerve fibre layer; VF-MD: visual field mean defect.

|  | **POAG**<br>**(N=20)** | **Controls**<br>**(N=15)** | ***p*-value** |
| --- | --- | --- | --- |
| **Demographic data and comorbidity** |  |  |  |
| Average age (y) | 64.85 | 70.27 | 0.11 |
| Females (%) | 25 | 53.33 | 0.09 |
| Mean BMI (kg/m<sup>2</sup>) | 25.75 | 28.27 | 0.30 |
| Diabetes (%) | 25 | 13.33 | 0.39 |
| Hypertension (%) | 35 | 73.33 | 0.02 |
| Hyperlipidaemia (%) | 25 | 60 | 0.04 |
| Thyroid disease (%) | 5 | 0 | 0.33 |
| **Systemic medications** |  |  |  |
| Anti-hypertensives (%) | 35 | 73.33 | 0.02 |
| Lipid-lowering medications (%) | 25 | 60 | 0.04 |
| Antiplatelet therapy (%) | 25 | 13.33 | 0.39 |
| Oral diabetes medications (%) | 25 | 13.33 | 0.39 |
| Insulin (%) | 0 | 0 | - |
| Corticosteroids (%) | 5 | 0 | 0.33 |
| Thyroid hormone (%) | 5 | 0 | 0.33 |
| Oestrogen (%) | 0 | 0 | - |
| Vitamin D (%) | 10 | 6.67 | 0.73 |
| Others (%) | 40 | 33.33 | 0.69 |
| **Ophthalmological features and glaucoma medication** |  |  |  |
| Mean visual acuity (LogMar) | +0.05 | +0.03 | 0.37 |
| Mean IOP (mmHg) | 15.82 | 13.84 | <0.001 |
| Mean CCT (μm) | 544.44 | - | - |
| Average RNFL thickness (μm) | 68.7 | - | - |
| Mean VF-MD (dB),<br>(eye with worse MD) | -3.99 | - | - |
| Glaucoma severity (%) |  |  |  |
| Mild | 80 | - | - |
| Moderate | 10 | - | - |
| Severe | 10 | - | - |
| Glaucoma medications (%) |  |  |  |
| Beta-blockers | 60 | - | - |
| Prostaglandin analogue | 85 | - | - |
| Alpha-2-agonists | 0 | - | - |
| Carbonic anhydrase inhibitor | 15 | - | - |

### Plasma nicotinamide concentrations

The univariate analysis of the results obtained using the semi-quantitative LC-HRMS method on plasma samples from the initial cohort revealed significant differences between individuals with POAG and controls, with nicotinamide being the most discriminant metabolite (False Discovery Rate corrected *p* = 0.0027), showing an average nicotinamide decrease of 36 % in individuals with POAG compared to controls (Figure A).

**Figure:**
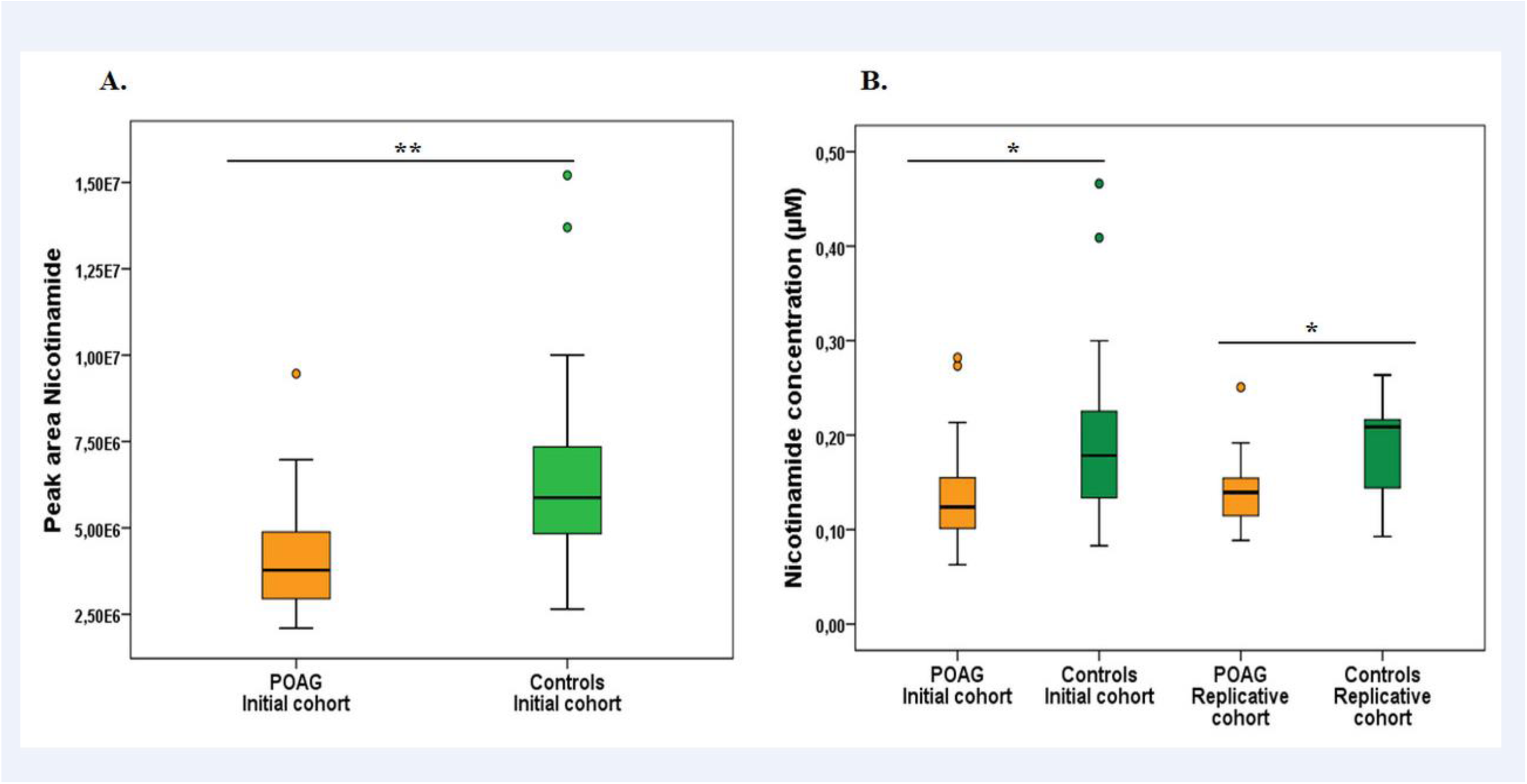
Boxplots showing nicotinamide levels in the initial (34 POAG and 30 control individuals) and replicative (20 POAG and 15 control individuals) cohorts. Error bars represent ± SEM, and the black bars within the boxplots represent the median concentration for each group. (A) Peak area of nicotinamide found in the initial cohort following LC-HRMS analysis discloses a glaucoma/controls fold change of 0.65. (B) Concentrations of nicotinamide found in the initial and replicative cohorts following LC-MS/MS analysis. The glaucoma/controls fold changes were 0.70 and 0.67 for the initial and replicative cohorts, respectively. The *p*-values between groups for all conditions were *: *p* <0.05 and **: *p* < 0.01.

This observation, subsequently tested in both the initial and replicative cohorts, using an independent quantitative measurement of nicotinamide designed for a clinical laboratory setting, supported the results obtained with the metabolomics analysis (Figure B). The median concentrations of nicotinamide found in individuals with POAG and controls were 0.12 µM (0.06-0.28 µM) *vs*. 0.18 µM (0.08-0.47 µM), and 0.14 µM (0.09-0.25 µM) *vs*. 0.21 µM (0.09-0.26 µM), respectively, in the initial and replicative cohorts, corresponding to a reduction of 30 % (*p* = 0.022) and of 33 % (*p* = 0.011) of the nicotinamide concentration in the initial and replicative POAG *vs.* control cohorts, respectively. The mean concentrations of nicotinamide found in individuals with POAG and controls were 0.14 µM *vs*. 0.19 µM, and 0.14 µM *vs*. 0.19 µM, respectively, in the initial and replicative cohorts.

During the semi-quantitative LC-HRMS several metabolites related to nicotinamide were assessed: 1-Methylnicotinamide, 6-hydroxy-nicotinic acid, nicotinic acid, nicotinamide mononucleotide, and NAD. Only 1-methylnicotinamide was accurately detected, but this metabolite was not discriminant between POAG and controls.

## DISCUSSION

Mitochondrial dysfunctions and decreased NAD content are hallmarks of aging in most organs ^13,14^ and many experimental studies, essentially performed on mouse models, have revealed that strategies based on NAD repletion effectively reverse age-related phenotypes and disorders ^15,16^, such as those affecting the skeletal muscles ^17^, the brain ^18^, and the endothelium ^19^. Recent studies on the DBA/2J mouse model of glaucoma, have further confirmed a dose-dependent protective effect of NAD repletion on the optic nerve, reaching a protection level of 93% at the highest nicotinamide dose tested (2000 mg/kg/day), despite a continuously elevated IOP ^6,7,20^. More importantly, the age-dependent vulnerability of the RGCs in these mice was correlated with the decreased concentration of NAD in the retina. Thus, the nicotinamide deficiency we observed in the blood of POAG individuals parallels the NAD depletion observed in the DBA/2J mouse model. Interestingly, our study of plasma samples from individuals affected by dominant optic atrophy due to OPA1 mutations, another form of an age-dependent progressive optic neuropathy due to mitochondrial impairment, also revealed a 50 % reduction of nicotinamide whose chemical formula is C_6_H_6_N_2_O ^21^.

The main function of NAD as a redox cofactor consists in providing electrons from oxidized nutrients to the mitochondrial respiratory chain complex I, thus sustaining ATP production. In parallel, NAD-consuming enzymes, such as those involved in DNA repair, e.g. poly (ADP-ribose) polymerase (PARP), may consume NAD stocks excessively during aging, in particular to prevent the accumulation of DNA mutations ^13^. This excessive NAD consumption may compromise NAD-dependent complex I activity, the deficiency of which is frequently associated with inherited optic neuropathies, because of the particularly high energy required by RGCs to transduce visual information from the retina to the brain. In this respect, lymphoblasts of patients with POAG showed a mitochondrial complex I deficiency reflecting a systemic mitochondrial impairment ^4,5^. In addition, using targeted metabolomics on the plasma of POAG patients compared to controls, we have recently shown a metabolic profile combining the impaired utilization of energetic substrates and decreased levels of polyamines, attesting a mitochondrial dysfunction, and premature ageing ^22^. Since nicotinamide is one of the main contributors to the regeneration of NAD through a salvage metabolic pathway, nicotinamide deficiency could reflect excessive age-related NAD consumption, which subsequently leads to complex I deficiency, and the energetic failure responsible for the degeneration of RGCs.

Despite extensive research in the literature, we were unable to find normative values for plasma nicotinamide levels in normal subjects. We believe that this can be explained by a technological gap, since the plasmatic nicotinamide levels are very low in humans. We assume that the recent technological advances in mass spectrometry have allowed us to perform these measures and we can only hope that further independent studies will explore this area.

The main limitation of this study consists in the relatively small number of individuals in both the initial and replicative cohorts. However, we found a significant decrease in vitamin B3 levels in patients with POAG compared to controls using two different techniques, with highly similar results in the two independent cohorts. Further studies with larger cohorts are also required, as well as investigations in populations with various cultural dietary habits, to find out whether this deficiency is consistently associated with POAG and eventually with other forms of glaucoma. Finally, the convergence between recent studies showing that oral administration of nicotinamide prevents glaucoma in the DBA/2J mouse model ^6,7,20^ and our study on patients with POAG, opens promising therapeutic perspectives based on nicotinamide supplementation.

## Abbreviations and acronyms

BMI: body mass index
CCT: central corneal thickness
CPP: comité de protection des personnes
HESI: heated electrospray ionization
HRMS: high resolution mass spectrometry
IOP: intraocular pressure
IS: internal standard
LC: liquid chromatography
MRM: Multiple Reaction Monitoring
NAD: Nicotinamide adenine dinucleotide
NM: Nicotinamide
NM-d_4_: nicotinamide-d_4_
OCT: optical coherence tomography
PFP: pentafluorophenyl
POAG: primary open-angle glaucoma
RGC: retinal ganglion cell
RNFL: retinal nerve fibre layer
VF-MD: visual field mean defect

## ACKNOWLEDGMENTS

We acknowledge support from the *Institut National de la Santé et de la Recherche Médicale* (*INSERM)*, the *Centre National de la Recherche Scientifique* (*CNRS)*, the *University of Angers*, the *University Hospital of Angers*. We also thank the following patients’ foundations for their support: “*Fondation VISIO*”, “*Ouvrir les Yeux*”, “*Union Nationale des Aveugles et Déficients Visuels*” “*Association contre les Maladies Mitochondriales*”, “*Retina France*”, “*Kjer France*”, “*Fondation Berthe Fouassier*”, *“Fondation pour la Recherche Médicale”* and *“Association Point de Mire”*.

We are grateful to Kanaya Malkani for critical reading and comments on the manuscript and to Dr. Odile Blanchet and the team of the *Centre de Ressources Biologiques* of the University Hospital of Angers for processing the biobank samplings.

