## Supplementary materials for "Nicotinamide Deficiency in Primary Open-Angle Glaucoma"

### Supplementary Material

#### Table of contents

Supplementary Figure…………….………………………………………………………………. S-2

Supplementary Table ....………….…………………………………………………………………. S-3

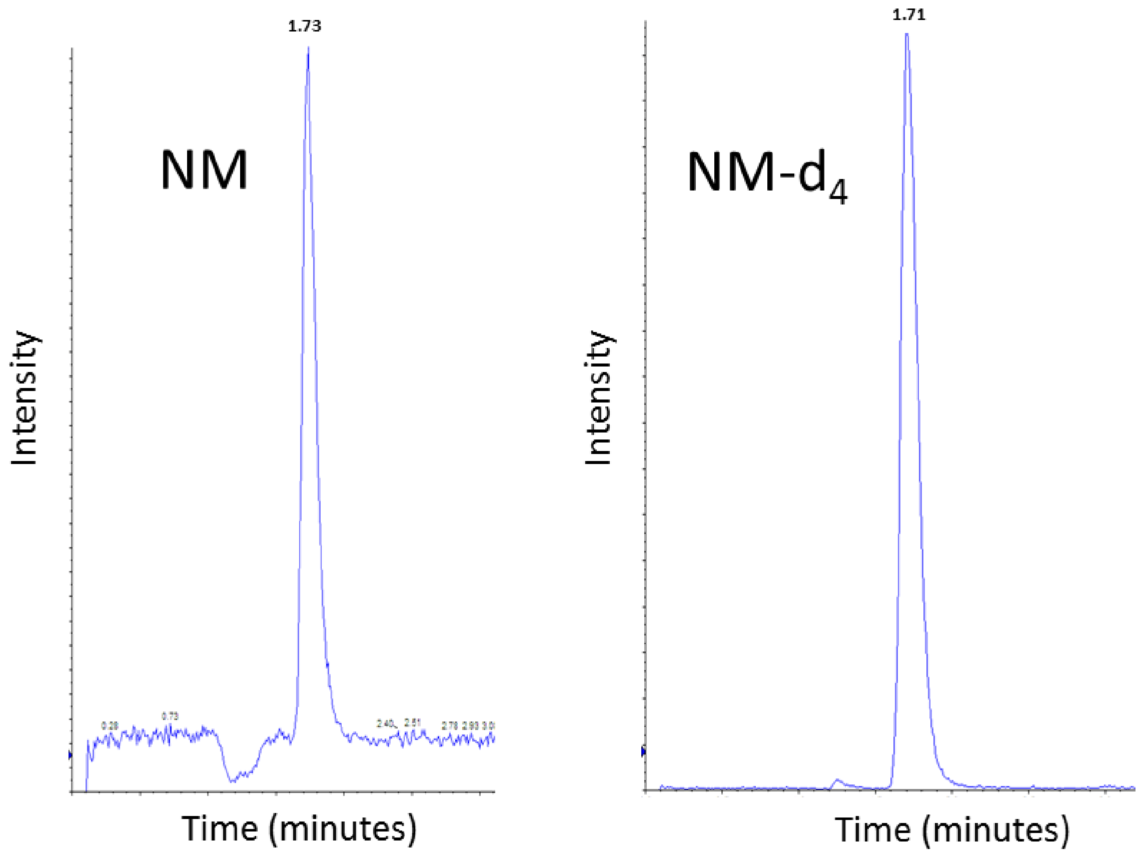


**Supplementary Figure: Chromatograms obtained for nicotinamide (NM; left) and its isotope-labelled analogue nicotinamide-d$_4$ (NM-d$_4$; right) in a plasma sample.** Deuterated nicotinamide, NM-d$_4$ was used as internal standard for the absolute quantification of nicotinamide concentration in the plasma from the cohorts. For quantitative analysis, the use of a stable isotope-labelled analogue of the analyte, as internal standard, is recommended to correct for possible loss during sample preparation steps and matrix effects, during mass spectrometry acquisitions. The ideal internal standard is a substance not contained in the sample, structurally related to the analyte (NM in our case), and having a retention time close to that of the analyte (retention times of NM and NM-d$_4$ are 1.73 min and 1.71 min, respectively). NM-d$_4$ perfectly fulfils these criteria.

**Supplementary Table:** The distribution of the blood collection hour between patients and controls.

| Time of sample collection | 8-12 am | 1-4 pm | *p*-value |
|---|---|---|---|
| POAG (initial cohort) | 18 | 16 | |
| Controls (initial cohort) | 16 | 14 | $p = 0.97$ |
| POAG (replicative cohort) | 12 | 8 | |
| Controls (replicative cohort) | 9 | 6 | $p = 1$ |
